# Population dynamics and entrainment of basal ganglia pacemakers are shaped by their dendritic arbors

**DOI:** 10.1101/388504

**Authors:** Lior Tiroshi, Joshua A. Goldberg

## Abstract

The theory of phase oscillators is an essential tool for understanding population dynamics of pacemaking neurons. GABAergic pacemakers in the substantia nigra pars reticulata (SNr), a main basal ganglia (BG) output nucleus, receive inputs from the direct and indirect pathways at distal and proximal regions of their dendritic arbors, respectively. We combine theory, optogenetic stimulation and electrophysiological experiments in acute brain slices to ask how dendritic properties impact the propensity of the various inputs, arriving at different locations along the dendrite, to recruit or entrain SNr pacemakers.

By combining cable theory with sinusoidally-modulated optogenetic activation of either proximal somatodendritic regions or the entire dendritic arbor of SNr neurons, we construct an analytical model that accurately fits the empirically measured somatic current response to inputs arising from illuminating various portions of the dendritic field. We show that the extent of the dendritic tree that is illuminated generates measurable and systematic differences in the pacemaker’s phase response curve (PRC), causing a shift in its peak. Finally we show that the divergent PRCs correctly predict differences in two major features of the collective dynamics of SNr neurons: the fidelity of population responses to sudden step-like changes in inputs; and the phase latency at which SNr neurons are entrained by rhythmic stimulation, which can occur in the BG under both physiological and pathophysiological conditions.

Our novel method generates measurable and physiologically meaningful spatial effects, and provides the first empirical demonstration of how the collective responses of SNr pacemakers are determined by the transmission properties of their dendrites. SNr dendrites may serve to delay distal striatal inputs so that they impinge on the spike initiation zone simultaneously with pallidal and subthalamic inputs in order to guarantee a fair competition between the influence of the monosynaptic direct- and polysynaptic indirect pathways.

**Author Summary:** The substantia nigra pars reticulata (SNr) is a main output nucleus of the basal ganglia (BG), where inputs from the competing direct and indirect pathways converge onto the same neurons. Interestingly, these inputs are differentially distributed with direct and indirect pathway projections arriving at distal and proximal regions of the dendritic arbor, respectively. We employ a novel method combining theory with electrophysiological experiments and optogenetics to study the distinct effects of inputs arriving at different locations along the dendrite.

Our approach represents a useful compromise between complexity and reduction in modelling. Our work addresses the question of high fidelity encoding of inputs by networks of neurons in the new context of pacemaking neurons, which are driven to fire by their intrinsic dynamics rather than by a network state. We provide the first empirical demonstration that dendritic delays can introduce latencies in the responses of a population of neurons that are commensurate with synaptic delays, suggesting a new role for SNr dendrites with implications for BG function.

## Introduction

The basal ganglia (BG) are a collection of forebrain nuclei involved in various aspects of motor control and habit formation. The substantia nigra pars reticulata (SNr) is one of the main output nuclei of the BG, innervating the ventral thalamus, superior colliculus and reticular formation [1–3]. SNr GABAergic neurons receive thousands of synaptic inputs. Most of them are inhibitory inputs arising from direct pathway spiny projection neurons (dSPNs) in the striatum [4,5], or from the external segment of the globus pallidus (GPe) in the indirect pathway [6,7]. Excitatory inputs originate from the subthalamic nucleus (STN) [8,9], the pedunculpontine nuclei (PPN) [10], and, to a lesser extent, from the cerebral cortex [11]. Thus, in the SNr, inputs from the direct and indirect pathways converge onto the same neuron.

The vast majority of SNr GABAergic neurons are autonomously active and discharge continuously and regularly at 6-30 spikes/s in rodents *in vitro* [12–15], even when their synaptic inputs are completely blocked and at higher rates *in vivo* [16–18]. A rhythmically firing neuron – also called a pacemaker neuron – can be represented by a phase variable that advances from 0 to 1 as the somatic voltage of the pacemaker neuron advances from one action potential threshold to the next. Such a phase oscillator is sensitive to the timing of impinging inputs. Thus, in the case of a pacemaker neuron, each individual synaptic input is sufficiently small so that its only lasting effect is a finite change in the phase of the pacemaker that either postpones or advances the next spike. The sign and amplitude of this phase change depend on the phase of the pacemaker’s cycle at which the synaptic input arrived. Thus, the complex neuronal dynamics of synaptic integration by the pacemaker are reduced to phase change as a function of the timing of inputs [19–22]. This reduction vastly simplifies the treatment of the collective dynamics of populations of pacemakers.

The quantity that describes the susceptibility of the pacemaker neuron’s phase to small voltage perturbation as a function of the timing of the perturbation is called the neuron’s phase response curve (PRC) [19–22]. Although the phase reduction and the use of PRCs are theoretically valid only for weak perturbations, they prove to be very useful in practice. The PRC in response to somatic stimulations can be easily measured experimentally, and has been used successfully to predict neurons’ responses to arbitrary patterns of somatic current injections [23–30]. Thus, the phase reduction is computationally simple, can capture the essence of the neuronal dynamics, and can be readily implemented experimentally on real neurons.

The use of phase reductions and PRCs has been applied both experimentally and theoretically [23,24,27,29,30], by and large, only to somatic voltage perturbations. However, neurons can possess elaborate dendritic arbors. Previous theoretical and experimental work has suggested that PRCs need to be redefined and generalized to accommodate inputs arriving at the dendritic arbors [31–37]. These studies predicted that taking the dendritic localization of inputs into consideration could have a measurable effect on collective neurons dynamics. Here, we put this prediction to test for the first time, and do so for actual neuronal pacemakers with dendrites.

SNr neurons in rodents possess dendrites that can extend up to 0.75 mm long, usually branching only once or twice [38,39]. Excitatory inputs from the STN are distributed along the entire length of the dendrites [40,41]. In contrast, the two major inhibitory inputs are differentially distributed: inputs from the GPe, belonging to the indirect pathway, are located at the soma and on proximal dendrites [42–45], while direct pathway striatal synapses impinge upon smaller distal dendrites and terminal tufts [45]. The functional relevance of the differential spatial organization of direct and indirect pathway inputs is unknown.

We combine theory and electrophysiological experiments in acute brain slices from transgenic mice that express channelrhodopsin-2 (Thy1-ChR2 mice) in GABAergic SNr neurons [46,47] to investigate the impact of dendritic localization of synaptic inputs on the control of spike timing in these pacemakers. This assay allows us to directly perturb the membrane voltage at various locations on the somatodendritic tree. In order to compare stimulation of proximal *vs.* distal locations on the dendritic arbor, we illuminate either a small region containing the soma and proximal dendrites or a wider field of view containing the entire dendritic tree, while measuring somatic currents and voltage. This approach does not take into account the complex morphology of the cells. However, eliminating the effect of the variability arising from the shape of individual dendritic arbors to hopefully generate more robust results. Moreover, it allows us to use a simple cable theoretic model, in which the entire dendritic tree is collapsed to a single cable, to fit the experimental results [48].

We begin by investigating the electrotonic length of SNr GABAergic pacemakers using periodic perturbations, and constructing a model that accurately predicts the somatic current response to inputs arising from stimulating various portions of the dendritic arbor. Next, we show that the choice of whether to illuminate only the soma and proximal dendrites or the entire dendritic arbor generates measurable and systematic differences in the PRCs corresponding to each of these conditions. Finally, we show that the divergent PRCs correctly predict differences in two major features of the collective dynamics of SNr neurons. The fidelity and latency of population responses to sudden changes in inputs, on the one hand, and the phase latency at which SNr neurons are entrained by rhythmic stimulation, on the other, are both determined by the extent of the dendritic tree illuminated – in close agreement with predictions arising from their respective empirical PRCs.

This is the first experimental demonstration of the impact of the electrotonic length of the dendritic arbor on the collective dynamics of pacemaker neurons. Our results suggest a possible role for the differential spatial distribution of inputs onto SNr GABAergic neurons, where SNr dendrites may serve to delay distal striatal inputs so that they impinge on the spike initiation zone simultaneously with pallidal and subthalamic inputs. Coherent rhythmic inputs to SNr neurons arise under various physiological (e.g., sleep, [49]) and pathophysiological (e.g., parkinsonism, [18,50]) conditions. Thus, the way in which these neurons follow their inputs, or become entrained to them, will depend on the spatial distribution of their inputs throughout their dendritic arbor (e.g., distal striatal vs. proximal pallidal inputs).

## Results

### A cable theoretic model of optogenetic dendritic activation

In order to investigate the impact of the dendritic location of inputs on the SNr pacemakers, we employed our previously published model of a semi-infinite cable attached to a pacemaking axosomatic compartment [32]. Active dendritic conductances are incorporated into the model by linearizing them in the vicinity of the dendrite’s resting membrane potential, resulting in a quasi-linear cable *u(x,t)* (Fig. 1A). The cable’s Green function (or impulse response) describes the spatio-temporal voltage response to a brief perturbation applied at the origin at time *t* = 0 (with vanishing boundary conditions at ± ∞) and provides a method to solve the cable’s partial differential equation. The Green function is given by

**Figure 1:**
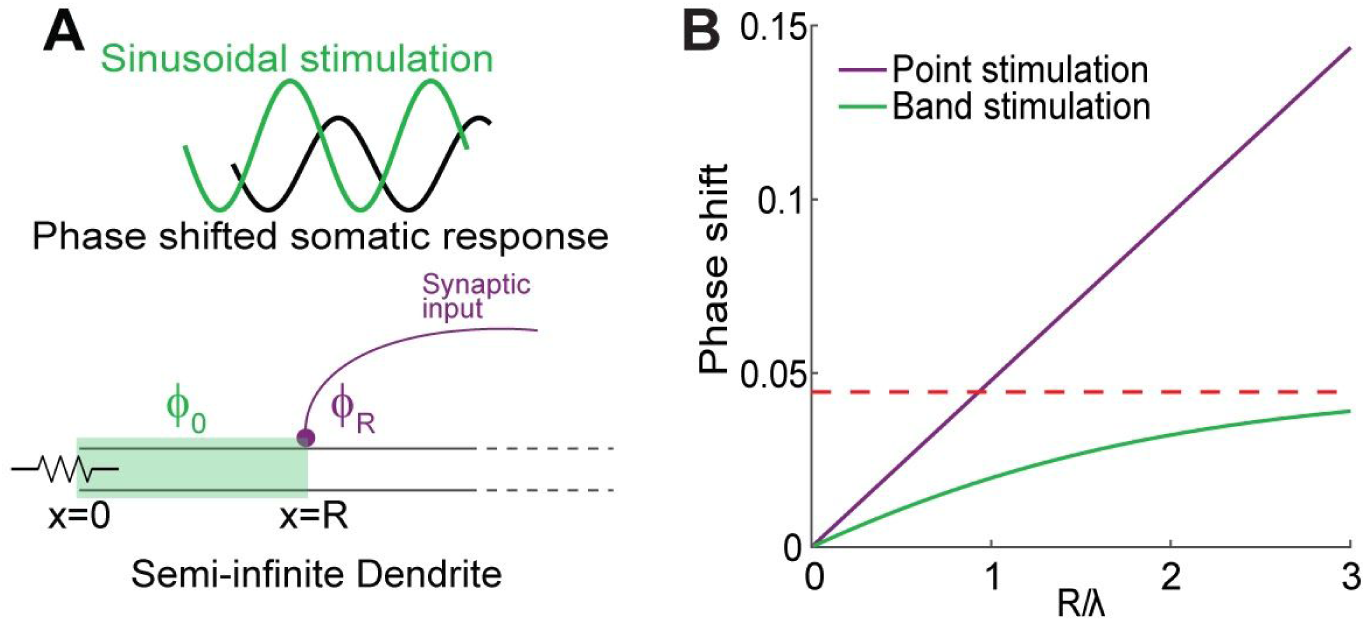
A cable theoretic model of optogenetic dendritic activation. **A:** Our model of the dendrite consists of a semi-infinite linear cable, where the somatic boundary at *x=0* is kept at a constant holding potential *u(0,t)=u*_*0*_. Optogenetic stimulation is modeled as a band of illumination starting at the soma and ending at *R* (green). Alternatively, *x=R* can be the locus of a point illumination (purple line indicates afferent axon terminating at the point). Stimulus is a temporal sine waveform (green) and the somatic current response is an attenuated and phase shifted version of it (black). The phase shift of the somatic response is Φ_0_ for point illumination and Φ_*R*_ for band illumination starting at *x=0* and ending at *x=R*. **B:** The phase shift between a sinusoidal optogenetic stimulation and the corresponding somatic current response as a function of the ratio between *R* and the dendrite’s electrotonic distance λ for band (green) and point stimulation (purple). The phase shift for band stimulation is bounded (red), while point stimulation is not bounded. Both curves are calculated for *f=10 Hz* and τ = 10 *ms*.

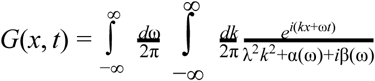

Where *λ* is the characteristic length of the cable, *k* is the wave number and ω is the angular frequency in the spectral representation of *G*. *α(ω)* and *β(ω)* are the contributions from the passive cable properties as well as terms that arise from linearizing the nonlinear conductances in the cable [32,33]. In particular, for a linear cable α(ω) = 1 and β(ω) = ωτ [32], where τ is the time constant of the cable.

Because we are interested in the current injected into the soma by the cable, we assume that it is semi-infinite where the somatic boundary at *x* = 0 is kept at a constant holding potential *u*(0, *t*) = *u*_0_ forall *t.* In the voltage clamp experiments this assumption is enforced experimentally. Thus, the current injected into the soma is proportional to the spatial derivative of *u*(*x*, *t*) at *x* = 0, with a proportionality factor κ:

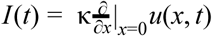

The Green’s function for the semi-infinite (si) cable with these boundary conditions is given by

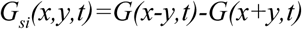

and describes how a voltage perturbation at location *y* propagates to location *x*.

The optogenetic stimulation of the cell is modeled as a band of illumination that starts at distance *r* from the soma and ends at distance *R*, where the stimulus has the temporal waveform *cos*(ω *t*), with ω = 2π*f* where *f* is the driving frequency. To calculate the current injected into the soma it is necessary to integrate the input from *y=r* to *y=R* and over all time. Thus,

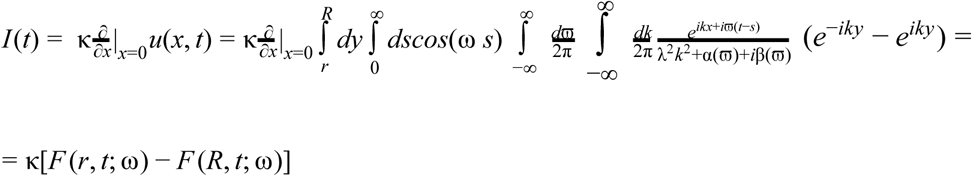

Where

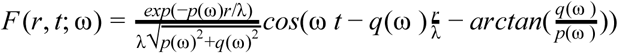

With

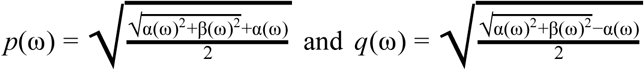

This means that illuminating a band of the cable with a sinusoidal temporal envelope generates a somatic current injection with two sinusoidal contributions, each with an amplitude that decays exponentially with distance of the boundaries of the band from the soma, and each with two contributions to the phase shift relative to the phase of the input. One shift that scales with the distance and another that does not. The spatial decay and all phase shifts increase with the driving frequency *f*.

To measure the resulting phase shift between the injected current *cos*(ω *t*) and the somatic current response *I(t)*, we calculate the location of the peak of the cross-correlation function (CCF) between them by averaging their product over all time. The CCF is given by:

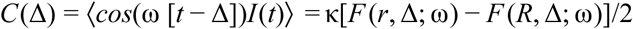

Differentiating with respect to Δ and comparing to zero results in the following equation:

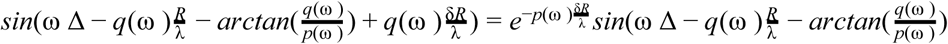

where 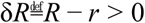. This equation can be used to find the phase shift Φ = ω δ corresponding to the peak in the CCF. We consider two cases:

*Case I - the stimulated region is small compared to the electrotonic length (*δ*R*≪λ):

In this case, after linearization we get the equation

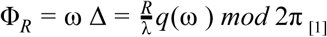

At this limit, the phase behaves like the stimulation of a single point at distance *R* from the soma (Fig. 1A, purple dot).

*Case II - the stimulated region begins at the soma (r* = 0):

In this case (Fig. 1A, green band), we get

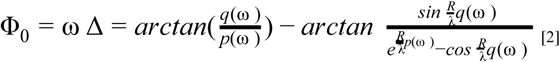

To summarize, the model allows us to calculate the current injected by the cable into the soma in response to a sinusoidal current perturbation with frequency *f*, given at some location *R* in the dendrite (case I), or applied to a region of the dendrite beginning at the soma (case II). Our analysis shows that the current arriving at the soma is also sinusoidal, but it lags by a phase delay of Φ_*R*_/2π or Φ_0_/2π, respectively.

Note that the phase delay following a field stimulation is bounded in the case of a linear dendrite [*e.g.*,α(ω) = 1 and β(ω) = ωτ], as the distance from the soma grows (*R* → ∞, eq. 2). In contrast, the phase shift in response to point stimulation (eq. 1), continues to increase linearly with the distance from the soma (Fig. 1B).

### Investigating the electrotonic length of SNr dendrites

To measure the effect of the dendritic localization of inputs on the somatic current response, we used transgenic mice that express ChR2 throughout the entire somatodendritic arbor of SNr neurons [46,47]. We chose to compare two extreme cases: illuminating either the soma and proximal dendrites (a diameter of ~ 130 μ*m* around the soma, see Materials and Methods) or the entire dendritic tree. This choice was motivated by three considerations. First, we reasoned that the strategy of illuminating the entire region is likely to create a more consistent and robust effect across diverse geometries of dendritic arbors. Second, the cable theoretic model we use to fit the data is more applicable under these conditions, as illuminating a band of cable in the model is comparable to illuminating an annulus in the plane, centered around the soma. Note that in our model the edges of the stimulated region are sharp, while our empirical stimulation scheme generates some falloff between the center of the illuminated area and its boundaries. However, the transition region is small compared to the difference between the diameters corresponding to the two conditions, adding a negligible error to the estimate generated by our model. Finally, both situations could conceivably be physiologically relevant. Coherent rhythmic input could arise under physiological conditions such as sleep [16,49] and pathophysiological conditions, such as parkinsonism [18,50]. Coherent pallidal input would preferentially target the soma and proximal dendrites [42,43,45], whereas coherent subthalamic input could activate the entire dendritic arbor [40,41].

To ensure that our proximal stimulation condition indeed activates a portion of the dendritic arbor, we employed two-photon laser scanning microscopy and measured the diameters of SNr somata. GABAergic SNr cells tend to be elongated, and the diameters of their somata measured 17.7 − 33.74 μ*m* (with a mean of 24.99 ± SD of 4.88 μ*m*, n=15) along the larger axis. Thus, both stimulation conditions activate dendrites as well as the cell’s soma and they differ in the extent of the dendritic arbor that they stimulate.

Hence, we optogenetically stimulated the ChR2-expressing SNr neurons using 470 nm LED light that was sinusoidally modulated at various temporal frequencies (0.25 to 16 Hz, 3-4 seconds of stimulation per frequency), while blocking all glutamatergic and GABAergic inputs. The cell was held at –70 mV to prevent spiking, and the current injected to the soma was measured in whole-cell voltage clamp. This was repeated under the two stimulation conditions: a) proximal stimulation targeting the soma and proximal dendrites; and b) full-field illumination exciting the soma as well as the entire dendritic arbor (Fig. 2A).

**Figure 2:**
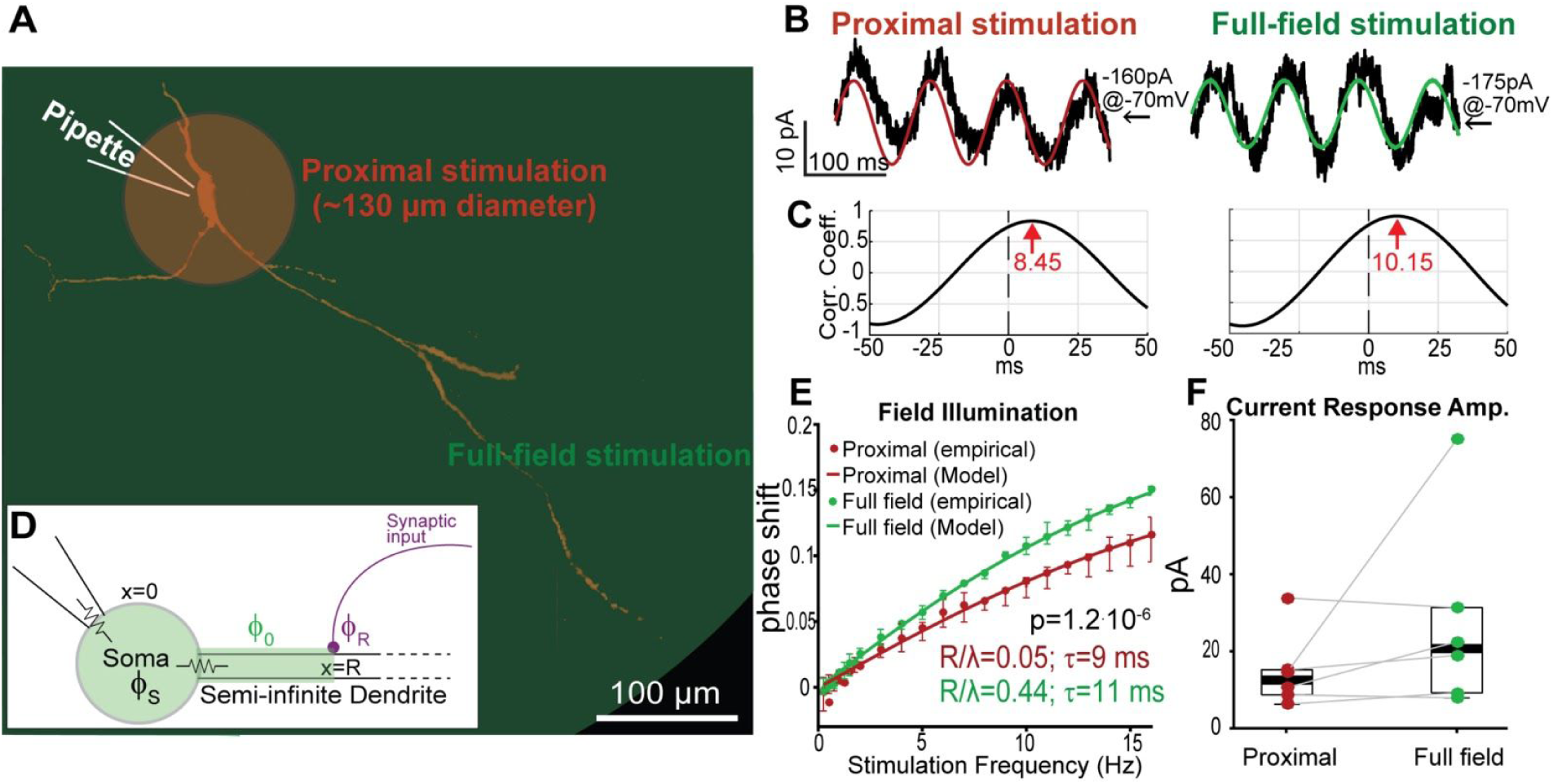
Somatic current responses to periodic illuminations of SNr GABAergic neurons shift depending on the size of the dendritic arbor activated. **A:** Collapsed two-photon image of typical SNr GABAergic neuron, filled with Alexa Flour. To compare distal and proximal inputs, optogenetic stimulation was applied under two conditions: 1) illuminating a diameter of ~130 μ*m* around the soma, targeting the soma and proximal dendrites (proximal, red); and 2) full-field illumination stimulating the entire dendritic arbor (full-field illustration). **B:** Optogenetic sinusoidal waveforms at various frequencies (0.25-16 Hz) were delivered under the proximal (red) and full-field (green) stimulation conditions. Somatic whole-cell current response (black) is a phase shifted sinusoid. **C:** Phase shift between optogenetic periodic stimulation and somatic current response was determined by the location of the peak in the cross-correlation function between the two traces (red arrows). **D:** The model used to fit the data consists of a semi-infinite cable, as in figure 2A, that is connected to an isopotential soma which adds an additional angular phase Φ_*S*_. **E:** Proximal and full-field illumination conditions give rise to two distinct curves (n=6; p=1.2.10^-6^, ANCOVA). The response to rhythmic stimulation shows a spatial effect – phase shift between stimulation and somatic response is larger under full-field illumination (green) compared to proximal (red). Under the assumption of a linear dendrite, we can fit the _0*S*_ empirical results (points) with our model ((Φ_0_ + Φ_*S*_)/2π, lines). Error bars represent standard error of the mean. **F:** Amplitude of somatic current response was comparable between the proximal and full-field stimulation conditions. Central marks indicate the median and box edges represent 25th and 75th percentiles.

The current measured with a patch electrode at the soma in response to the sinusoidal optogenetic stimulation was a phase shifted sinusoidal waveform of the same temporal frequency (Fig. 2B). The phase shift between the periodic stimulation and the somatic response was determined by the location of the peak in the CCF between the two traces (Fig. 2C). This was used to generate a plot of the phase shift as a function of stimulation frequency for each illumination condition (Fig. 2E). Amplitudes of somatic current responses were similar for proximal and full-field illumination (Fig. 2F), and we saw no relationship between current response amplitude and the magnitude of the phase shift. However, the two spatial conditions give rise to distinct curves (n=6; *p* = 1.2 · 10^−6^, ANCOVA). As expected, the phase increases with frequency in both conditions. The phase shift between the stimulation and the somatic current response is larger when the entire dendritic field is illuminated compared to only the soma and proximal dendrites (Fig. 2E). Thus, the response of SNr GABAergic neurons to subthreshold rhythmic stimulation shows a spatial effect, indicating that optogenetic methods can reveal the electrotonic length of the dendrites and their contribution to somatic currents. Importantly, the experiment was repeated with holding voltages of –60 mV and –50 mV and yielded similar results, indicating that there is no voltage dependence of the phase shifts in the subthreshold range (S1 Figure). In other words, under these stimulation conditions the dendrites do not exhibit substantial subthreshold nonlinearities.

In order to estimate the electrotonic properties of SNr dendrites, we fit the parameters τand 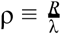 of our cable theoretic model to our data [48]. The empirical curves describing the dependence of the phase shift on frequency were fit to the appropriate case II of the model (eq. 2). Because the electrode does not measure the current injected by the dendrite into the soma directly, but rather the current at the patch pipette tip, we need to add to the dendritic phase shift (predicted by the model) the phase introduced by the somatic membrane (Fig. 2D). Assuming an isopotential passive membrane (which is a good approximation when holding at subthreshold voltages) yields the additional angular phase of Φ_*S*_= *arctan* (2π*f* τ) (Fig. 2D). Even with the simplifying assumption of a strictly passive linear dendrite – which is in line with the voltage independence of the phase shift we described above – we attain a good agreement between the predictions of our model and the empirically observed average frequency dependent phase shift at the soma (Fig. 2E). Fitting our theoretical curve to the empirical data for the two conditions results in similar and physiologically plausible [51] values for the membrane time constants (i.e., τ _*proximal*_ = 9*ms*, τ _*full*−*field*_ = 11*ms*, Table 1) and the characteristic length λ. The ratio ρ of the boundary of the stimulated region to the dendritic characteristic length 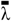 was also extracted (ρ*proximal* = 0.05, ρ*full*−*field* = 0.44 _*full*−*field*_, Table 1). From ρ*proximal* we can deduce that the effective characteristic length of the SNr dendrites is approximately 1.3 mm (=65 µm/0.05) [51]. Moreover, the model estimates that the value corresponding to illumination of the entire dendritic field is ~9 times larger than the value for illumination of only the soma and proximal dendrites. Therefore, we can deduce that on average dendrites are light-activated out to ~580 µm from the soma. This is a reasonable estimate, considering that in acute slices regions beyond that are often cut or are too deep to be affected by the illumination. The experiment repeated at −60 mV (S1 Figure) gave rise to a different estimate of ρ = 0.22, but because the estimates at −70 mV and −50 mV were similar we trusted these. If we were to choose an average of the three estimates it would lower our estimate of the diameter up to which the dendrites were activated but it would still be several hundred micrometers out. In summary, our theoretical and experimental results demonstrate that distal and proximal inputs undergo different phase shifts by the time they reach the soma. We therefore hypothesize that these different phase shifts will also differentially affect the suprathreshold behavior of these pacemaking neurons.

**Table 1:** *values of fitted parameters*

| *Section* | *Parameter* | *Proximal* | *Full-field* |
| --- | --- | --- | --- |
| *Electrotonic characteristics*<br>*@ -70 mV* | $\tau$ | *9 ms* | *11 ms* |
| | $\rho = \frac{R}{\lambda}$ | *0.05* | *0.44* |
| *Current response kinetics* | $\tau_{\text{rise}}$ | *$1.5 \pm 0.26 \text{ ms } (n=11)$* | *$3.6 \pm 0.81 \text{ ms } (n=11)$* |
| | $\tau_{\text{decay}}$ | *$5.1 \pm 1.2 \text{ ms } (n=11)$* | *$4.9 \pm 1.8 \text{ ms } (n=11)$* |
| *PRC estimation* | $\theta$ | *0.831* | *0.763* |
| | $A$ | *0.0103* | *0.0179* |
| | $C$ | *-0.0041* | *-0.0053* |
| *PSTH estimation* | $\theta$ | *0.901* | *0.733* |
| | $\sigma$ | *$5.713 \text{ (ms)}^{-\frac{1}{2}}$* | *$4.3704 \text{ (ms)}^{-\frac{1}{2}}$* |

### Proximal and full-field PRC estimation

To address this hypothesis, we estimated proximal and full-field PRCs by incorporating our two illumination conditions into a previously described method for PRC measurement using optogenetic pulses [47]. The neurons were subjected to Poisson-like processes composed of barrages 0.5-1 ms long light pulses, separated by exponentially distributed inter-pulse intervals (Fig. 3A, see Materials and Methods).

**Figure 3:**
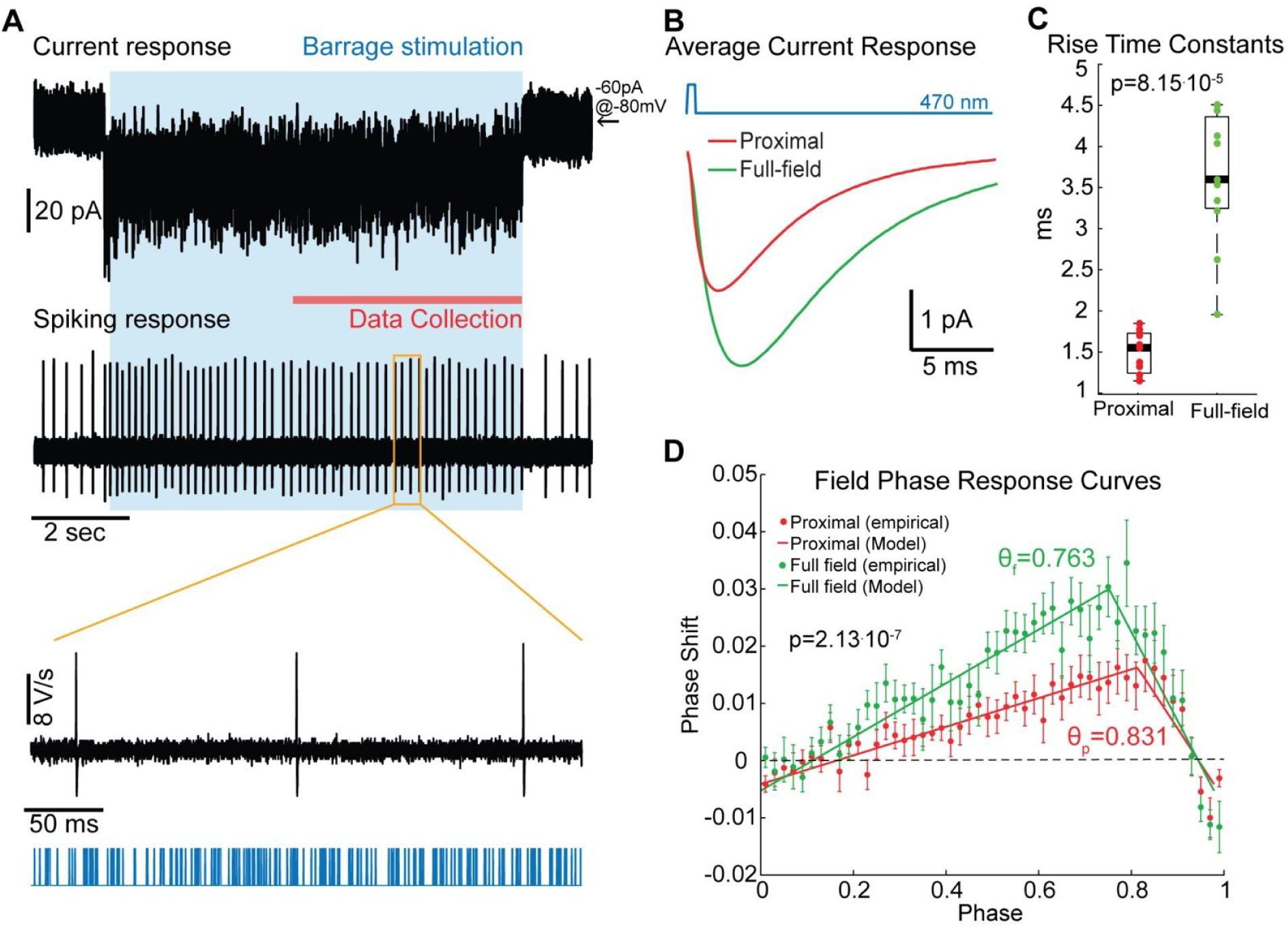
Proximal and full-field barrage stimulations generate distinct PRCs in SNr GABAergic pacemakers. **A:** ChR2 current and spiking responses (top, black) to barrage stimulation (bottom, blue). The stimulus generated an inward current in whole-cell voltage clamp and an increase in firing rate in cell-attached current clamp. Blue shading indicates stimulation time and red line marks the period in which ChR2 responses have reached steady-state and data is collected. **B:** ChR2 current responses to a light pulse (blue), averaged over all pulses and all cells (n=11). Full-field illumination (green) induces a longer response than proximal illumination (red). **C:** For each stimulation condition, a double exponential was fit to the average current responses for each cell and time constants were extracted (τ _*rise*_, τ _*decay*_). The distribution of rise time constants is significantly different in the two stimulation conditions (p=8.15.10^-5^, RST), and values are larger in the full-field condition compared to proximal. Central marks indicate the median and box edges represent 25th and 75th percentiles. **D:** Barrage stimulation was used to measure PRCs corresponding to the two stimulation conditions. Points are the empirical data and lines represent the fitted curves. Proximal and full-field illumination resulted in two distinct PRCs (n=19, p=2.1.10^-7^, ANCOVA), with the peak of the full-field curve occurring at an earlier phase than that of the proximal one (*θ*_*f*_=0.763 and *θ*_*p*_=0.831, respectively). Error bars represent standard error of the mean.

In order to examine the ChR2 currents generated by the barrage stimulation, cells were hyperpolarized to prevent spiking and steady-state currents (4-9 seconds after beginning of stimulation to avoid any effects of ChR2 deactivation) were measured in the whole-cell voltage clamp configuration (Fig. 3A). For each cell (n=11), a curve portraying the average current response to a light pulse was generated (Fig. 3B). As previously reported, average responses were well fit by a double exponential, chosen to accommodate ChR2 deactivation kinetics [47] (eq. 9, see Materials and Methods), and rise and decay time constants (τ _*rise*_, τ _*decay*_) were extracted (Table 1). Decay time constants were not significantly different for the two conditions (S2 Figure). However, rise time constants were significantly larger when the entire dendritic tree was illuminated compared to only the soma and proximal dendrites. This is evident in the distribution of rise time constants (*p* = 8.15 · 10^−5^, RST) (Fig. 3C), as well as the average current response curves for the different conditions (Fig. 3B). The delayed peak of the somatic response when the entire dendritic tree is illuminated is consistent with the delayed full-field response that we observed in the previous section. Thus, the barrage stimulation induces disparate somatic current profiles under the proximal and full-field stimulation conditions.

We have shown previously that the more distal a dendritic perturbation is, the more leftward a shift it will generate in the dendritic PRC compared to the somatic one [32]. Thus, the shift we observed in the current profile generated by the full-field illumination relative to the proximal one buttresses the hypothesis that the empirical PRC measured under the full-field illumination should exhibit a leftward shift relative to that measured under proximal illumination. To test this hypothesis, we applied the barrage stimulation to spontaneously firing SNr cells recorded in the cell-attached configuration (Fig. 3A) and estimated the two PRCs corresponding to the proximal and full-field illumination conditions. For each stimulation condition and for every cell, the PRC was estimated using a multiple linear regression method [47,52] (see Materials and Methods). Previous measurements of the PRCs of SNr GABAergic pacemakers reveal a triangular form - the PRC slopes upwards before peaking and falling to zero as the phase approaches 1 [47]. We therefore fit a triangle (parameterized by a parameter 0 < θ < 1 for the location of the peak, an amplitude *A* and an offset *C*) to our experimental data (see Materials and Methods), in order to extract the location of the peak (Table 1).

Our empirical data include negative values around phases 0 and 1 (Fig. 3D) due to jitter in the pacemaker’s unperturbed period in the course of the recording. Due to the refractory period of the action potential, phase theory dictates that the PRC should pass through the points (0, 0) and (0, 1). Thus, the empirical negative values are inconsequential. However, we focus on the phase of the PRC’s peak for the two stimulation conditions (θ_*proximal*_, θ_*full*−*field*_), and these values remain almost unchanged when the curve is forced to be zero at the beginning and end of the period.

Our analysis results in two distinct average PRCs (n=19; *p* = 2.13 · 10^−7^, ANCOVA), with the full-field PRC peaking before the proximal one (Fig. 3D). The steep negative slope at late phases represents the cells approaching the causal limit, where the input elicits an immediate spike [53]. According to our theoretical model, as well as the results of our study of the dendrite’s electrotonic length, the somatic current generated by a stimulation delivered in the full-field condition would be phase delayed compared to a proximal stimulation, which explains why the peak of the full-field PRC is shifted leftward relative to the proximal PRC.

Hence, differentially located inputs arrive at the soma at different times, thereby shifting the full-field PRC leftward relative to the proximal one. Next, we turn to measure properties of the collective activity of SNr pacemakers which theory predicts are dependent on the shape of the PRC [53–56]. First, we measure the fidelity and latency of the population response of SNr neurons to sudden changes in their input, and we then consider the propensity of rhythmic inputs to entrain SNr pacemaker neurons.

### Populations response to proximal vs. full-field stimulation

Barrage stimulation changed the firing frequency of SNr GABAergic cells (Fig. 4A). The perturbation significantly increased the pacemaker’s mean firing rate by 33% for proximal (n=19; *p* = 1.32 · 10^−4^,SRT) and by 19% for full-field (*p* = 1.5 · 10^−3^, SRT) illumination. Importantly, the distributions of baseline (*p* = 0.82, RST) and perturbed firing rates (*p* = 0.45, RST) were not significantly different across the two stimulation conditions (Fig. 4B). In a population of cells, the response to a step stimulation consists of two stages - an initial increase in firing and a relaxation to a steady-state rate. The rapidity at which the PSTH can track the change in inputs is a measure of the fidelity of the neurons’ coding of the input. The shape of the PSTH has been shown to be theoretically related to that of the PRC [56]. The Fokker-Planck formalism enables one to predict this relationship (See eq. 8 in Materials and Methods) [54,55]. Interestingly, this connection demonstrates that the initial rise in the PSTH should reflect the mirror image of the falling phase of the PRC [54]. Thus, the leftward shift in the full-field PRC should generate a rightward shift in the rise time of the full-field PSTH relative to the proximal PSTH. Estimation of the empirical PSTHs under both conditions confirmed this prediction, with the peak in the proximal PSTH appearing approximately 12 ms before that of the full-field PSTH (Fig. 4C). Moreover, fitting the empirical PSTH to the theoretical equation for the PSTH with an underlying triangular PRC, enables us to extract independent estimates of the peak of the triangular PRC based on the rise time of the PSTH (θ_*p*_ = 0.901, θ_*f*_ = 0.733, Table 1). Indeed, this estimation gave rise to a larger (but comparable) estimate of the phase shift between the peaks of the underlying proximal and full-field PRCs. A caveat in PRC estimation is the difficulty to estimate it empirically near the causal limit [53]. Thus, for weak perturbations, values of PRC peaks extracted from the PSTHs may provide a more precise estimation of the true shape of PRC’s late downswing, which may be occluded by the causal limit.

**Figure 4:**
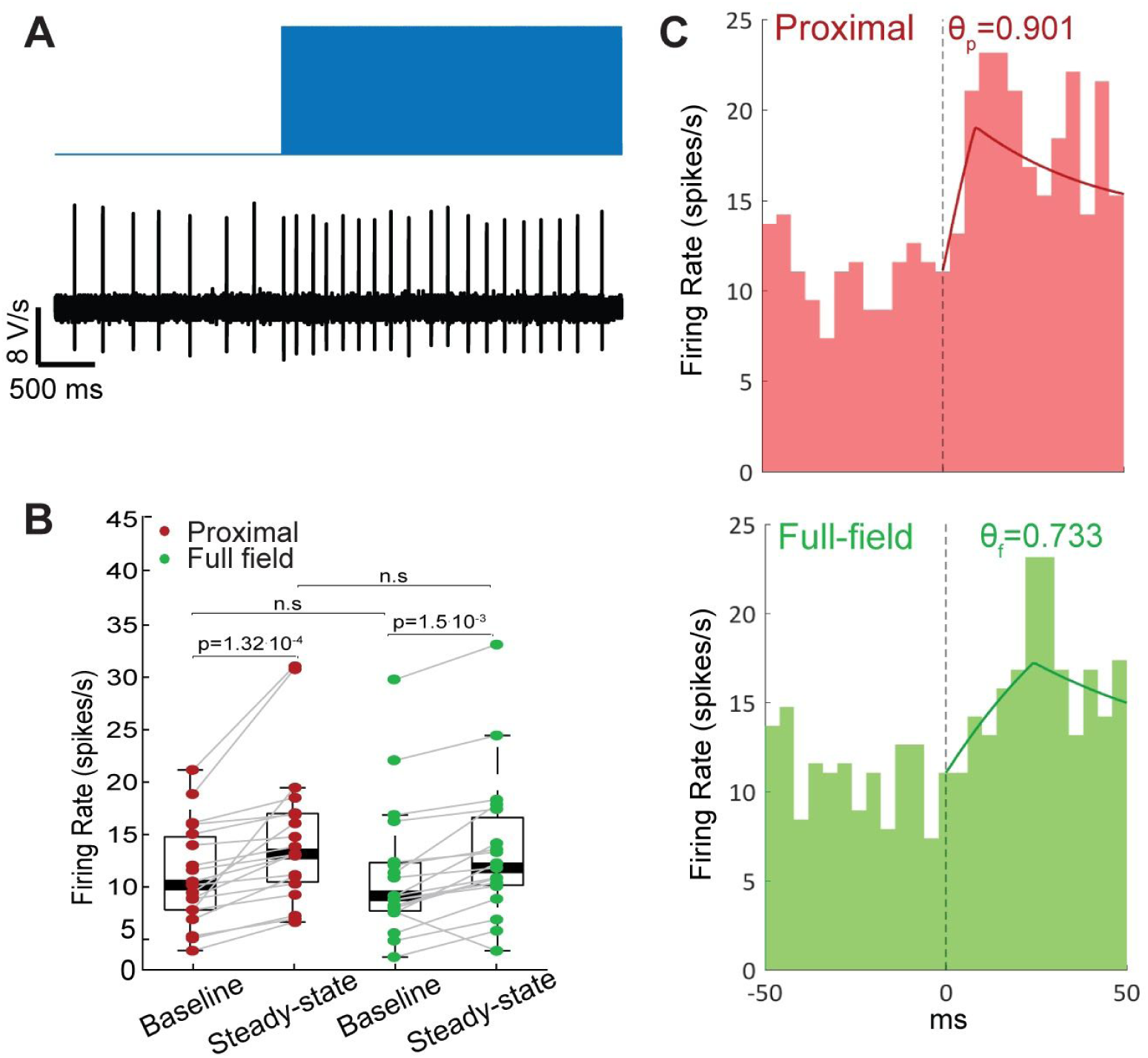
Proximal and full-field barrage stimulations induce distinct peri-stimulus time histograms (PSTHs). **A:** Example of an increase in the cell’s firing rate in response to barrage stimulation. **B:** Baseline and steady-state firing rates were similar under the proximal (red) and full-field (green) stimulation conditions (n=19). Barrage stimulation increased the firing rate significantly (proximal: *p* = 1.32 · 10^−4^; full-field: *p* = 1.5 · 10^−3^; SRT). Central marks indicate the median firing rates and box edges represent 25th and 75th percentiles. n.s - not significant. **C:** Peristimulus time histograms (n=19 cells, 25 repetitions per cell) centered around stimulus onset. The peak in population firing rate response to stimulus onset is delayed by approximately 12 ms in the full-field illumination condition (green) compared to the proximal illumination condition (red). Lines represent fitted theoretical curves, corresponding to θ_f_=0.733 and θ_p_=0.901.

### Entrainment of SNr pacemakers to rhythmic inputs

The discharge of a regularly active neuron can be entrained by small amplitude sinusoidal currents delivered to the soma, when the driving frequency is close to the neuron’s natural firing rate (or to integer multiples of it). Within this range of frequencies, the rhythmic stimulation reshapes the neuron’s firing pattern - the firing rate changes to match that of the periodic stimulation and becomes phase locked to it [57]. SNr neurons were allowed to fire spontaneously and their spiking activity was recorded in the perforated patch current clamp configuration. The perforated patch configuration was chosen to allow us to see subthreshold responses and injected somatic currents, while avoiding the disruption of pacemaking activity by the disruption of the neurons intracellular milieu.

A 30-60 seconds long 10-20 pA cosine shaped current of frequency *f* was injected into the soma of spontaneously firing cells (example in Fig. 5A). We studied the effect of the oscillatory input on the cell’s firing pattern based on a previously published method [57]. For each spike, we measured the duration of the next perturbed period, denoted *T*_*p*_, of the neuron’s spiking as a function of its effective phase, the phase Ψ in the period of the sinusoidal input in which the spike occurred, and fit a periodic function to the effective phase comprised of Fourier 3 modes, resulting in the function *T*_*p*_(Ψ) (Fig. 5B). Given an initial action potential at the effective phase Ψ_*n*_, we can predict the effective phase Ψ_*n*+1_ of the next action potential using the following iterative map:

**Figure 5:**
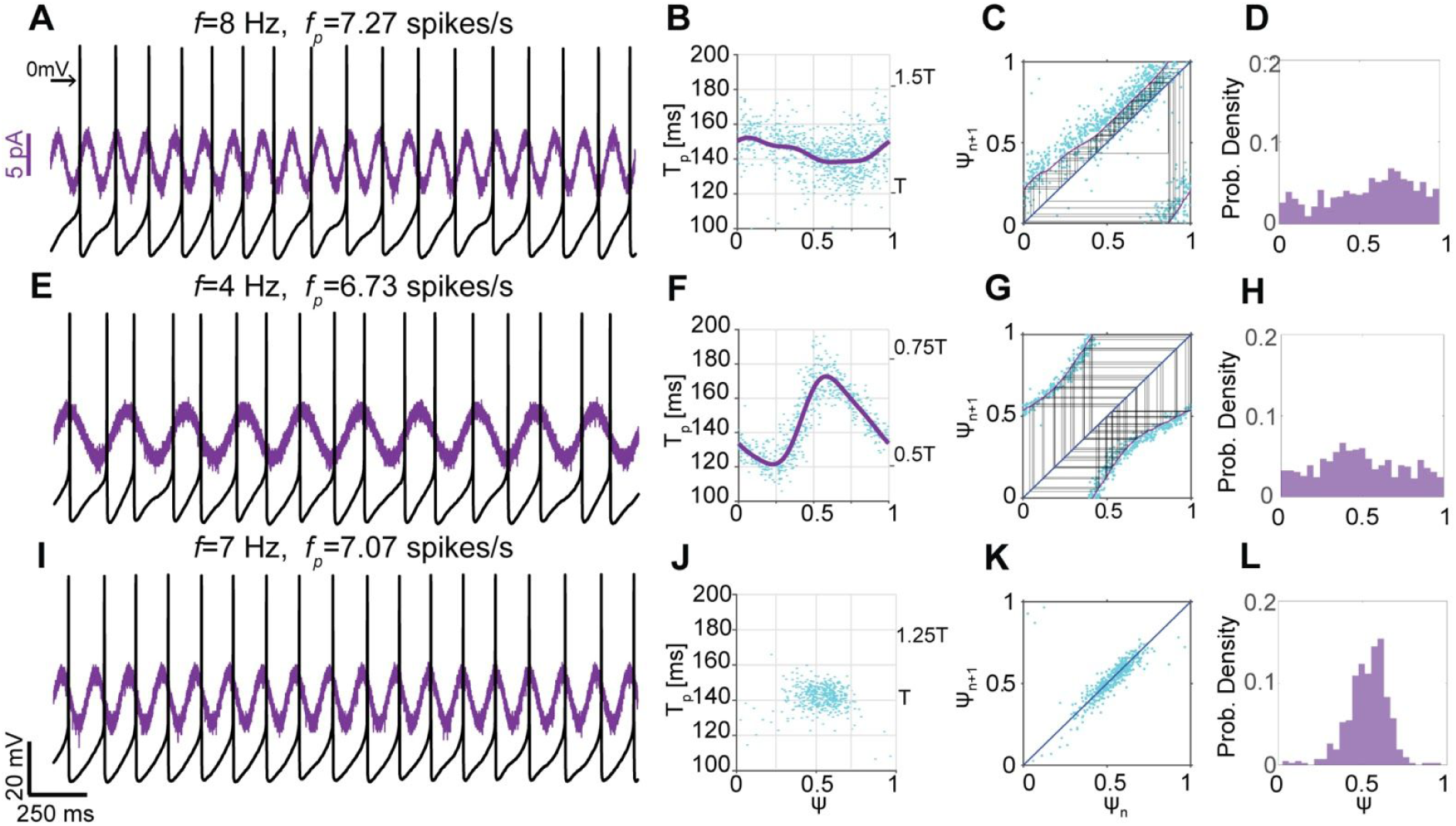
Entrainment of the spiking of SNr GABAergic pacemakers using sinusoidal somatic current injection. **A:** Perforated patch recording of a spontaneously firing cell with an average perturbed firing rate *f*_*p*_, as a cosine shaped current of frequency *f* was injected into the soma. When *f* is sufficiently larger than the natural firing rate *f*_*0*_, the pacemaker’s firing is not phase-locked to the periodic input. Left arrow indicates 0 mV. **B:** Scatter plot of the perturbed period of the cell’s spiking *T*_*p*_ as a function of the effective phase *Ψ* in which the spike occurred (cyan). Fitted curve (purple) was used to generate the iterative map shown in C. Right-hand-side y-scale shows *T*_*p*_ in units of *T*, the period of the rhythmic stimulus. **C:** An iterative map of the effective phase of the next spike *Ψ*_*n+1*_ versus that of the current spike *Ψ*_*n*_ (purple). Dots represent empirical data, and the map is generated from the fitted curve in B. The map is mostly above the diagonal and does not transect it (blue). Cobweb plot (black) uncovers chaotic dynamics. **D:** The distribution of effective phases is broad and samples the entire range between 0 and 1. **E, F:** Same as A, B with *f<f*_*0*_. **G:** Same as C with *f<f*_*0*_. The map (purple) is mostly below the diagonal (blue) and does not transect it. **H:** Same as D with *f<f*_*0*_. **I:** When *f~f*_*0*_, the cell’s spiking is phase-locked to the oscillatory input. The spike tends to occur at or slightly after the peak of the stimulus. **J:** Due to phase locking, the cell visits a narrow range of effective phases. All points of the perturbed period versus the effective phase are gathered around the equivalent phases 0 and 1. The period of the cell’s spiking is approximately constant and equal to the period of the stimulus *T*. **K:** All phase points are located on the diagonal and around phases 0 and 1, indicating phase locking. Therefore, a map cannot be estimated. **L:** The distribution of effective phases shows a clear peak around the phase of entrainment (~0.6).

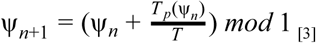

Therefore, the evolution of the phase of spiking relative to the oscillatory input can be represented using an iterative map. An intersection between the map and the diagonal (where Ψ_*n*+1_ = Ψ_*n*_) represents a fixed point of the dynamics. When the slope of the iterative map at the fixed point is smaller than unity, the fixed point is stable and the neuron is phase-locked with the stimulus. Thus, the sequence of phases visited by the neuron over a series of spikes can be predicted via this iterative map [57–59].

The fitted curve for the measurements of *T*_*p*_ as a function of Ψ (Fig. 5B) was used to generate an iterative map of effective phases (eq. 3). Note that the mapping closely traces the empirical scatter plot of Ψ _*n*+1_ versus Ψ_*n*_ (Fig. 5C). In the example presented in figure 5, the cell was spiking spontaneously at a frequency *f* _0_ ≈ 7 spikes/s throughout the recording. When the stimulation frequency *f* is larger than the cell’s natural firing frequency *f*_*0*_ (Fig. 5C), the map passes mostly above the diagonal and does not transect it, which is consistent with the lack of phase locking in the raw data (Fig 5A). The lack of phase locking is evident in the chaotic dynamics exhibited by the trajectory in the iterative map of effective phases (Fig. 5C), as well as in the wide probability distribution of effective phases covering all phases between 0 and 1 (Fig. 5D). When *f* < *f*_*0*_ (Fig. 5E), the map passes below the diagonal. Again, the neuron’s firing is not entrained by the periodic stimulation and the effective phase mapping gives rise to chaotic dynamics (Fig. 5G). Nevertheless, the distributions of phases in both cases (Fig. 5D,H) show an accumulation of phases in the area where the map passes closest to the diagonal (Fig. 5C,G), demonstrating the effect of the sinusoidal input.

When *f* ~ *f*_*0*_, the firing of the cell is phase-locked to the stimulus at a phase close to 0.5 (Fig. 5I). In this case, the pacemaker only visits a limited range of effective phases. Therefore, we cannot fit a curve to *T*_*p*_ measurements (Fig. 5J) and an iterative map (eq. 3) cannot be estimated. Nevertheless, the entrainment of the neuron’s firing to the periodic input is clear from the narrow distribution of points in the scatter plot of Ψ _*n*+1_ versus Ψ_*n*_ and their accumulation on the diagonal (Fig. 5K), as well as the sharp peaks around the equivalent phases 0 and 1 in the probability distribution of effective phases (Fig. 5L).

Next, we incorporate optogenetic excitation into this scheme. The experiment above was repeated, but the temporal cosine waveform stimulation was applied optogenetically using the two spatial conditions of proximal (Fig. 6A) and full-field (Fig. 6E) illumination. Depending on the ratio between the stimulation frequency *f* and the natural firing frequency of the cell *f*_*0*_, the stimulus either entrained the spiking activity of the cell, or was ineffective in causing entrainment. The efficacy in phase locking also differed somewhat between the two conditions, with full-field illumination inducing entrainment more readily. In order to examine the dendritic effect on entrainment, we focused on instances where phase locking was obtained.

**Figure 6:**
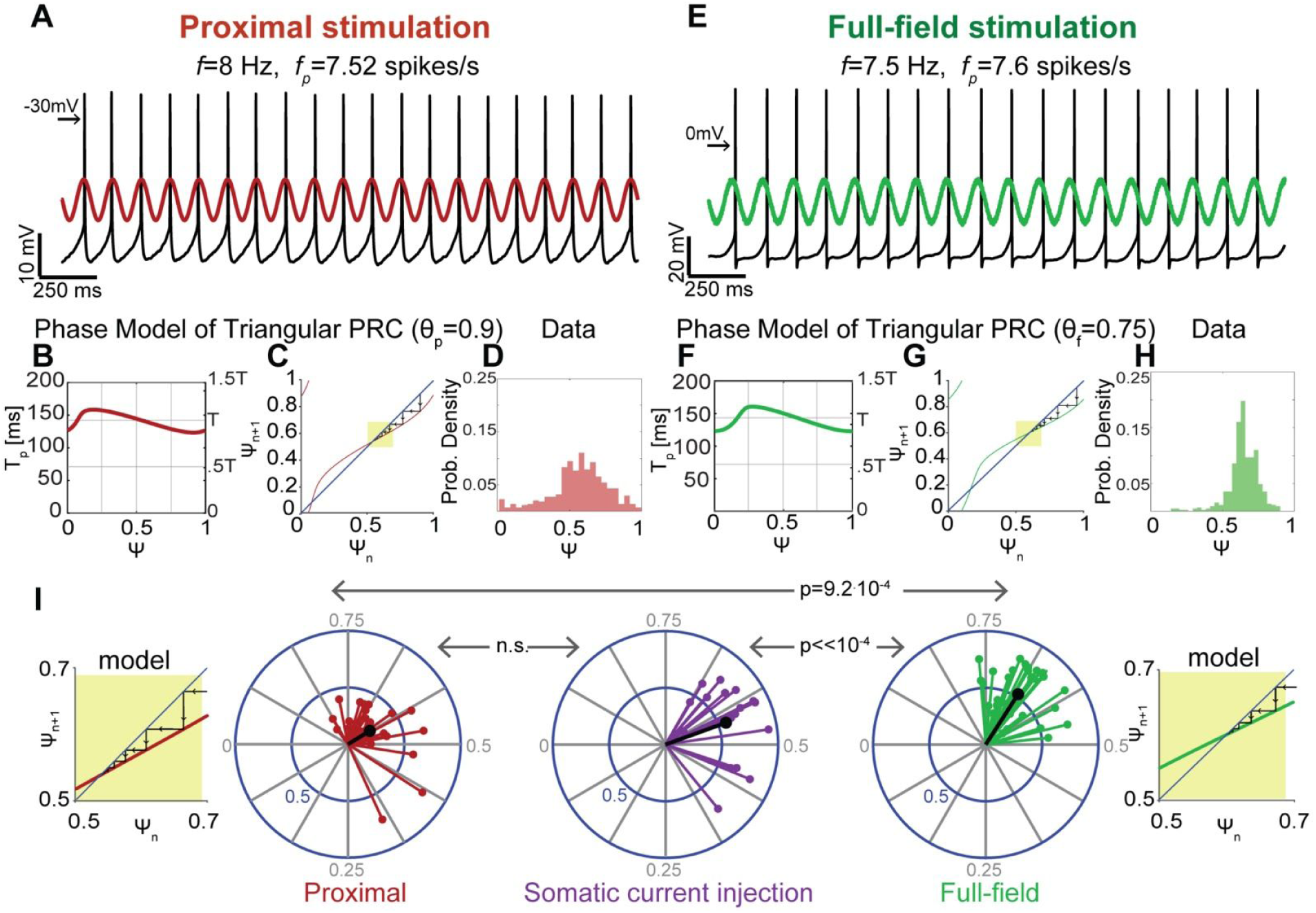
The dendrite impacts the phase of entrainment to rhythmic inputs. **A:** Perforated patch recording of a spontaneously spiking neuron with an average perturbed firing rate *f*_*p*_. **A** cosine shaped stimulation delivered in the proximal illumination condition entrained the discharge of the cell. The spike tends to occur near the peak of the stimulation, corresponding to phases near 0.5. Left arrow indicates −30 mV. **B:** Simulated (eq. 5) plot of the perturbed period (*T*_*p*_) as a function of the effective phase of the preceding spike (for θ _*p*_ = 0.9). **C:** Mapping of *Ψ*_*n+1*_ versus *Ψ*_*n*_ generated based on the simulated plot shown in B. The mapping (red) crosses the diagonal (blue), achieving a stable fixed point at phase 0.537, as demonstrated by the Cobweb plot (black arrows). Yellow box highlights stable fixed point. **D:** The probability distribution of effective phases shows a peak at early phases, in accordance with the raw data shown in A. **E:** Same as A for full-field illumination. Spikes occur slightly after peak of cosine stimulation. Left arrow indicates 0 mV. **F:** Same as B for full-field illumination (for θ_*f*_ = 0.75). **G:** Same as C for full-field illumination. The map (green) crosses the diagonal (blue), generating a stable fixed point at phase 0.6. Phase locking induced by full-field stimulation occurs at a delayed phase compared to proximal stimulation (shown in A-D). **H:** Same as D for full-field illumination. The phase of the peak in the probability distribution is later than that corresponding to proximal illumination and shown in D. **I:** Distribution of circular variances of effective phases (each calculated from a corresponding probability distribution of effective phase as in panels D and H) for various stimulation frequencies (2.5-21Hz) in n=6 cells, under three stimulation conditions: proximal illumination (red), somatic current injection (purple) and full-field illumination (green). Mean vectors are shown in black. Proximal illumination and somatic current injection induce locking in a similar range of phases, but current injection is more effective in entraining the neuron’s spiking. Phase locking generated by full-field illumination is shifted rightward compared to proximal illumination and current injection by 0.572-0.604. Yellow insets are zoomed-in versions of panels C and G that show that the shift in the model of the empirical distribution of locked phases agrees with the phase model simulations. See Materials and Methods for derivation of P-values.

To obtain a deeper understanding of the effect of the portion of the dendritic arbor being activated on entrainment, we performed a simulation. When phase locking occurs, the pacemaker samples a narrow range of effective phases around the phase of entrainment, and it is thus impossible to estimate an *empirical* curve relating the perturbed period to the effective phase. However, the perturbed period can be described using the following *theoretical* equation

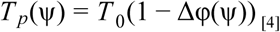

Where *T* _0_ is the unperturbed period of the oscillating neuron, and δΦ is the change in the intrinsic phase of the neuron induced by the stimulus. The evolution of intrinsic phase in response to a cosine shaped stimulation depends on the pacemaker’s PRC, denoted by *Z*, and is given by

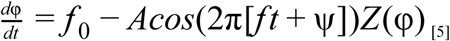

We evaluated the change in intrinsic phase δΦ by numerically integrating eq. 5 until the intrinsic phase Φ reaches 1 and measuring the resulting change δΦ for various values of the effective phase Ψ. PRCs were modeled as triangles peaking at θ_*p*_ = 0.9 and θ_*f*_ = 0.75 (values similar to those extracted from PSTH estimates) for proximal and full-field illumination, respectively. Both *f* and *f* _0_ were set to 7 Hz (*A* = 5). Simulated δΦ values were plugged into eq. 4, generating a *numerical* curve of *T*_*p*_ as a function of Ψ predicted by the phase dynamics and the PRC (Fig. 6B,F). This curve was then used to evaluate a mapping for the evolution of effective phases (eq. 3) under each illumination condition (Fig. 6C,G).

Both simulated mappings intersect with the diagonal achieving stable fixed points, which represent the effective phases of locking between the neuron’s spiking activity and the oscillatory stimulation. When a large portion of the dendritic field is stimulated, phase locking is predicted by the phase dynamics to occur at the phase 0.6 (Fig. 6G). In contrast, the predicted phase of locking corresponding to proximal stimulation is 0.537 (Fig. 6C). Thus, our simulation predicts that the effective phase of entrainment will occur slightly after the peak of the driving sinusoid and that this locking phase will be delayed when the entire dendritic arbor is stimulated compared to when only the soma and proximal dendrites are stimulated. This is consistent with the relative locations of the peaks in the probability distributions of empirical effective phases under the two spatial conditions (Fig. 6D,H).

This experiment was repeated using various stimulation frequencies (2.5-21Hz) in several cells (n=6). Oscillatory stimulation was delivered either optogenetically using proximal or full-field illumination, or as somatic current injection. For each trial, a probability distribution of effective phases was generated (Figs. 6D,H), and was used to calculate a circular variance vector [60] (see Materials and Methods). The argument of this complex vector represents the phase of locking between firing activity and the periodic stimulus, and its amplitude expresses the strength of locking. Because we are interested in instances of entrainment, only trials with circular variance amplitudes above a threshold determined by bootstrapping were included in the analysis (see Materials and Methods). While somatic current injection was more effective at inducing entrainment than proximal illumination (Fig. 6I), both stimulation conditions generated similar ranges of locking phases (averaging at 0.555 and 0.587 for current injection and proximal illumination, respectively). This suggests that the proximal stimulation condition, in which only the soma and proximal dendrites are activated optogenetically, is comparable to somatic current injections. In contrast, the typical phase of locking in the full-field illumination condition (averaging at 0.6585) is significantly delayed compared to the proximal illumination (*p* = 9.2 · 10^−4^, see Materials and Methods) or somatic current injections (*p* << 10^−4^, see Materials and Methods). Thus, the effective phase of the entrainment (shortly after the peak of the stimulation) and the difference between locking phases corresponds to those predicted by our simulation (Fig. 6C.G), which was based on the empirical PRCs for the two illumination conditions (Fig. 3D). The divergence of the PRCs, in turn, is explained by the electrotonic properties of the dendrites of the SNr GABA neurons (Figs. 2 & 3).

## Discussion

### Predictive models of dendritic neurons

To capture the entire dynamical repertoire of a neuron requires a detailed and high-dimensional model [61–63]. However, pacemaker neurons (and their corresponding models) offer an opportunity for simplification. Pacemakers traverse a low-dimensional closed contour called a limit cycle, so the high-dimensionality of the individual neuron can usually be shed in favor of a simpler dynamical system that captures the essence of the limit-cycle. The phase reduction method is a powerful formalism that does precisely that [19–22,64,65]. It reduces the pacemaker to a phase oscillator represented by its intrinsic frequency and its PRC, while maintaining the robustness and universality of pacemakers and oscillators. Pacemakers are attractive because the core of their dynamical properties *can* be measured and quantified in a single experiment. Many studies have capitalized on this and successfully measured empirical PRCs in order to characterize neuronal pacemakers and their collective dynamics [22–24,27,29,30].

However, reducing pacemakers to a phase oscillator with a single PRC, overlooks the contribution of their morphology to the phase dynamics. Previous studies investigated pacemaking using elaborate models that take into account detailed morphologies and ionic currents [34–37]. In the current study, we worked within the formalism of phase reduction theory, but still preserved the essential contribution of the dendritic arbor to the pacemaker’s dynamics. We combined previous results from cable theory [32,48,51], phase reduction theory [19–22,64,65], statistical physics [55,65,66] and nonlinear dynamical theory [57,59] to elaborate how the spatial extent of dendritic activation dictates the structure of the PRC and how that in turn characterizes the effect of dendrites on the population rate response and entrainment of pacemaker neurons. We aimed to generate the simplest model that would provide a plausible translation of the dendrite’s electrotonic length to temporal delays. While more detailed models can certainly describe our empirical data, remarkably, the very simple cable model in combination with these powerful and generic theories gave rise to predictions that were then corroborated experimentally.

The advantage of the phase reduction formalism is that it reduces high-dimensional oscillators to a one-dimensional limit-cycle represented by a single phase variable. This formalism does not need to be abandoned once a complex dendritic structure is included. Rather, with a relatively simple extension of the formalism developed for point neurons, one can add the effects of dendrites. This may be valuable for neuroscientists who study and want to simulate large populations of neurons. With this formalism they can relatively easily add the main impact of the dendrite due to its effective dendritic length, which, as we show is experimentally accessible.

### Interrogating electrotonic length of SNr neurons with sinusoidal optogenetic inputs

We began our combined theoretical and experimental study by investigating the electrotonic length of SNr GABAergic pacemakers, and generating a model that accurately fits the somatic current response to inputs arising from illumination of various portions of the dendritic field. Although our formalism can detect dendritic nonlinearities, our analysis showed that SNr GABAergic pacemakers and their dendrites are mostly linear under our stimulation conditions. This is supported by the good fit we obtained between the empirical data and the model under the assumption of a linear dendrite (Fig. 2E), as well as the fact that the effect was not dependent upon holding potential (S1 Figure).

Our novel method generated measurable and physiologically meaningful spatial effects. Fitting the model to the data yielded an estimate of electrotonic length of SNr dendrites of 1.3 mm, which is of the correct order of magnitude and consistent with the diameter of the dendrites [51]. However, our model is a vast simplification of the dendritic arbor and does not take into account the morphologies of individual cells. Moreover, this simplified model provides predictions regarding localized perturbations that could not be tested using our optogenetic stimulation method, which consisted of illuminating an entire portion of the dendritic field, beginning at and including the soma. The spatial effects that we detect using this method are robust but small. Interestingly, our theoretical model of the dendrite predicts that the spatial effect induced by a localized input would be significantly more dramatic (Fig. 1B). Taken together, this calls for a further investigation of the effect of the dendritic electrotonic structure using localized stimulation, which takes into account the structure of each specific dendritic tree.

The stimulation conditions that we employ could also be physiologically relevant. Because GPe targets the proximal somatic region [42,43,51], while the STN targets the entire dendritic tree uniformly [40,41], the proximal illumination could reflect a coherent activation of afferent GPe inputs, while the full-field illumination could reflect the coherent activation of afferent STN inputs. Coherent rhythmic input could impinge on SNr neurons both under physiological conditions, such as sleep [49] or anesthesia [16,18], and pathophysiological conditions, such as parkinsonism [50]. Thus, the spatial effect that we demonstrate could provide insight into how rhythmic pallidal and STN inputs may interact to drive and entrain SNr neurons, and possibly drive pathological oscillatory activity in Parkinson’s disease (PD) [66]. In this pathological context, it may be important to consider that the oscillatory inputs from the GPe and STN to SNr may maintain a specific phase relationship because they are reciprocally connected [9,67,68]. An interesting future direction would thus be to further examine the physiological significance of our findings by optogenetically activating natural afferents to SNr cells arising from the GPe, the STN and the striatum both *in vitro* and *in vivo.*

### Reduced single neuron dynamics and population responses

A fundamental question in the field of neuroscience is how the properties of the single cell manifest themselves in a network of neurons. Both theoretical and experimental work have demonstrated that the large scale network dynamics of neurons often depend on a simple dynamical property or characterization of the individual neurons, rather than their high-dimensional and morphologically complex description. For example, a large body of work, conducted over the past decade by Wolf and collaborators, has shown that the high-fidelity of the population rate response of cortical neurons arises from and depends on the rapidity of action potential initiation. Because cortical neuronal dynamics are governed by a balance between excitation and inhibition that keeps these neurons near threshold, when an abrupt change occurs in their shared input, sufficiently many neurons can cross threshold and follow the input. Therefore, the only rate limiting factor, in these fluctuation driven neurons, is how fast each one can generate an action potential [55,69,70].

This mechanism of stochastic resonance provides a good description of population responses in the cortex, the hippocampus and many other brain regions. However, in the BG, many neurons are not driven by noise but rather by their autonomous pacemaking. In this case, the framework for simplifying the description of neurons is the phase reduction method [19,21,64] where each neuron is characterized by its mean intrinsic rate and its PRC. Here too the high-fidelity of the population response can be derived from the PRC [54,55]. It turns out that the limiting factor on the fidelity of the response is the final descending part of the PRC. The initial rising phase of the PSTH, which is a reflection of the rapidity with which the population can respond, scales like the intrinsic period of the pacemaker and is roughly a mirror image of the final descending region of the PRC (see eq. 3). Here we provide an empirical demonstration of this previously known relationship, and show that a single global parameter - the effective electrotonic length of the dendrite 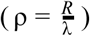 being stimulated, can capture measurable dendritic effects on the PRCs and PSTHs of SNr pacemakers.

Unlike in the case of fluctuation driven neurons, the pacemakers whose phase is closest to action potential threshold are the least responsive to input (the PRC vanishes there) and therefore cannot contribute to the population response. In order for populations of pacemakers to attain a high fidelity representation of their input they need to discharge faster. In primates and humans, SNr GABAergic cells exhibit very high discharge rates of up to 145 spikes/s [71], enabling SNr projection neurons to transmit information rapidly.

### Spiking resonances in SNr determine their entrainability

According to our analysis, SNr GABAergic cells and their dendrites are linear and do not exhibit subthreshold resonances. However, they do display spiking resonances - the firing of SNr pacemakers can be entrained to rhythmic inputs, if the stimulation frequency is close enough to the neuron’s intrinsic frequency – its natural discharge rate. We demonstrated this first using periodic somatic current injection. We then incorporated optogenetic stimulation into this scheme and demonstrated that the tendency of the SNr cells to be entrained to rhythmic inputs is significantly affected by the dendrite, with inputs arriving at different locations inducing distinct phases of locking.

The spiking resonance acts as a filter - signals containing frequencies near the natural discharge frequency of the neuron will be transmitted more effectively [57,72,73]. It is therefore possible that deep brain stimulation is only therapeutically effective at frequencies in the 120-140 Hz range [74], because it needs to successfully entrain BG output neurons (e.g., SNr) whose spiking resonances are in that range due to these pacemakers’ exceptionally high intrinsic firing rates [71].

The three stimulation methods that we employed - somatic current injection, proximal and full-field illumination, were not equally effective in entraining SNr neurons. Comparing the effects of current injection and optogenetic stimulation is complicated, as the two methods perturb the cell in inherently different ways. We deal with this issue by defining a bootstrapping based threshold (see Materials and Methods) and only considering instances where entrainment did occur, focusing on the dendritic effect on the phase of locking rather than the efficacy in entraining the cell. Interestingly, somatic current injection and proximal illumination induced similar phases of locking, while current injection was more similar to full-field illumination in the potency to entrain the spiking of SNr projection neurons. This strengthens the view that the two features - entrainment efficacy and phase of locking - are independent.

### Impact of dendritic structure on population responses

A recent theoretical study has argued that the impedance load of the dendritic arbor should affect the rapidness of spike initiation [75], which should invariably impact the response fidelity of fluctuation driven neurons such as pyramidal neurons. Similarly, but in the case of pacemakers, our study demonstrated that dendrites affect two aspects of the collective dynamics of SNr GABAergic pacemakers. First, we examined the fidelity of the population rate response and showed that the peak in the response induced by a stimulation of the entire dendritic arbor is delayed compared to stimulation of the soma and proximal dendrites. Next, we showed that when the firing of an autonomously active neuron is entrained by a periodic input, locking tends to occur at a later effective phase for activation of a larger portion of the dendritic arbor. Importantly, our study of the dendritic impact on currents arriving at the soma allowed us to relate – for the first time empirically – these population effects to the transmission properties of SNr dendrites. Because SNr neurons are actively decorrelated by recurrent connections [49,76] they are perfectly fit to function as a population readout of the integrated activity of the direct and indirect pathways. Thus, the rapidity and fidelity of transmission in SNr projection neurons are highly important.

### Implications of SNr dendritic delays on the race between go and no-go signals

The intracellular response latency of SNr GABAergic cells is typically attributed to synaptic delays [5,12]. However, our results suggest that the delay in the response of a population of neurons can originate from additional sources. We show, as previously suggested [54,56], that the fidelity of response in a population of oscillating neurons is determined by the shape of their PRCs, and that the PRC is in turn affected by the dendrite. The earlier peak in the response to proximal inputs, implies that inputs arising from the GPe, which impinge upon the soma and proximal dendrites, would generate a faster response than inputs originating from striatal spiny projection neurons and arriving at distal dendrites and terminal tufts. STN projections onto SNr neurons are distributed along the entire length of the dendrite, and would thus induce a delay that is longer than that generated by proximal GPe inputs, but considerably shorter than the delay in response to distal striatal inputs. The spatial organization of inputs can therefore have a significant effect on the latency in the response of a population of SNr neurons.

SNr GABAergic projection neurons integrate inputs from the direct and indirect BG pathways, with afferents arising from striatal dSPNs, the GPe and the STN converging onto the same cell. GABAergic inputs from direct pathway dSPNs are thought to promote movement by inhibiting SNr neurons, while glutamatergic inputs transmitted by indirect pathway STN neurons block behavior by exciting the same neurons. GABAergic GPe inputs are inhibited by indirect pathway spiny projection neurons resulting in the disinhibition of SNr cells and the further depressing of movement. Thus, an input originating in the striatum and activating both pathways simultaneously generates a competition between ‘go’ and ‘no-go’ signals. However, because the ‘go’ signal is a monosynaptic input whereas the ‘no-go’ signal is a polysynaptic input, the ‘go’ signal would have an inherent advantage in the race to inhibit or entrain SNr neurons. Here we show that dendritic delays are manifested in the PSTH rise time in SNr pacemakers, and thus have a direct impact on the relative timing between the two competing signals. This may act as a compensation mechanism, effectively delaying direct pathway inputs originating from the striatum so that they impinge on the axosomatic region of the SNr neuron together with the indirect pathway inputs that coursed through the polysynaptic route via the STN or GPe, enabling a fair competition between ‘go’ and ‘no-go’ signals.

A similar race is believed to underlie ea core function of BG circuits - the decision to interrupt and cancel actions [77–79]. According to this view, the successful interruption of an action depends on the outcome of a competition between a dSPN ‘go’ signal and a PPN induced STN ‘stop’ signal. Evidence of this race was observed in the PSTHs of SNr and STN neurons, and suggests that the fast glutamatergic STN signal will fail in inhibiting the action, if the slower to rise GABAergic dSPN ‘go’ signal arrives at the SNr early enough to shunt away its effects [17]. This race occurs at a significantly slower time scale than the dendritic delays that we detect. However, a fast SNr response to STN inputs is critical for effective action inhibition. Our findings suggest that the spatial organization of inputs onto SNr neurons gives the uniformly distributed STN inputs an advantage over distal striatal inputs in quickly affecting SNr projection neurons. Conversely, because the ‘go’ signal arising from dSPNs in the race model peaks in the SNr PSTH at approximately 100 ms after the ‘stop’ cue, it does not require SNr neurons to respond particularly fast which makes the distal location of dSPN synapses appropriate [17].

## Materials and Methods

### Animals

This study was carried out in accordance with the recommendations of and approved by the Hebrew University Animal Care and Use Committee. Experiments were conducted with 3-12-week old male and female homozygous transgenic Thy1-ChR2 mice [B6.Cg-Tg (Thy1-COP4/EYFP) 18Gfng/1]. These mice express ChR2 under the Thy1 promoter [46] in SNr GABAergic neurons. In SNr GABAergic cells, ChR2 is expressed in the soma as well as all parts of the dendritic field [47].

### Slice preparation

Mice were deeply anesthetized with ketamine (200 mg/kg)–xylazine (23.32 mg/kg) and perfused transcardially with ice-cold-modified artificial cerebrospinal fluid (ACSF) bubbled with 95% O2–5% CO2, and containing (in mM) 2.5 KCl, 26 NaHCO3, 1.25 Na2HPO4, 0.5 CaCl2, 10 MgSO4, 0.4 ascorbic acid, 10 glucose and 210 sucrose. The brain was removed, and 240 µm thick sagittal slices containing the SNr were cut in ice-cold-modified ACSF. Slices were then submerged in ACSF, bubbled with 95% O2–5% CO2, containing (in mM) 2.5 KCl, 126 NaCl, 26 NaHCO3, 1.25 Na2HPO4, 2 CaCl2, 2 MgSO4 and 10 glucose, and stored at room temperature for at least 1 h prior to recording.

### Electrophysiological recording

The slices were transferred to the recording chamber mounted on a Zeiss Axioskop fixed-stage microscope and perfused with oxygenated ACSF at 31 °C. To guarantee that the effects we measured were generated post-synaptically, the ACSF solution contained 10 μ*M* 6,7-Dinitroquinoxaline-2,3-dione (DNQX) to block AMPA receptors, 50 μ*M* D-(-)-2-Amino-5-phosphonopentanoic acid (D-AP5) to block NMDA receptors, 10 μ*M* 6-Imino-3-(4-methoxyphenyl)-1(6*H*)-pyridazinebutanoic acid hydrobromide (SR) to block GABA_A_ receptors, and 2 μ*M* (2*S*)-3-[[(1*S*)-1-(3,4-Dichlorophenyl)ethyl]amino-2-hydroxypropyl] (phenylmethyl) phosphinic acid hydrochloride (CGP) to block GABA_B_ receptors. An Olympus 60X, 0.9 NA water-immersion objective with a 26.5 mm field number (FN) was used to examine the slice using standard infrared differential interference contrast video microscopy. Patch pipette resistance was typically 4-5 MΩ when filled with recording solutions. The junction potential estimated at 7-8 mV was not corrected. For both whole-cell and cell-attached recordings the intracellular solution contained (in mM) 135.5 KCH3SO3, 5 KCl, 2.5 NaCl, 5 Na-phosphocreatine, 10 HEPES, 0.2 EGTA, 0.21 Na2GTP, and 2 Mg1.5ATP, pH 7.3 with KOH (280–290 mOsm/kg). For perforated patch recordings 2 μ*g*/*ml* of gramicidin B was added to the intracellular solution. In whole-cell voltage clamp recordings, cells were held between −70 to −80mV to avoid spiking, or at −50mV and −60mV to search for possible voltage dependencies in the subthreshold range. In cell-attached and perforated patch current clamp recordings, neurons discharged spontaneously, and figures depict the temporal derivative of the voltage. Electrophysiological recordings were obtained with a MultiClamp 700B amplifier (Molecular Devices, Sunnyvale, CA). Signals were filtered at 10 kHz online, digitized at 20 kHz and logged onto a personal computer with the Signal 6 software (Cambridge Electronic Design, Cambridge, UK).

### Optogenetic stimulation

Optogenetic stimulation was achieved with blue-light (470 nm) LED illumination via the objective (Mightex, Toronto, ON, Canada). Field illumination was applied under two conditions, in all experiments conducted in this study: a) proximal stimulation illuminating a ~130 μ*m* diameter around the soma (achieved by placing an opaque disk with a central pinhole at the 60X objective’s back focal plane), thereby targeting the soma and proximal dendrites (proximal); and b) full-field illumination stimulating the entire SNr (and beyond) with a 5X objective which excites the soma and the entire dendritic field (full-field, see S3 Figure). Given the small size of SNr GABAergic cell somata reported in the Results, both illumination conditions include dendritic activation. In all experiments, LED light intensity at the back plane of the objective was chosen such that stimulation generated comparable current and voltage responses for the two conditions. Because the opaque disk with the pinhole blocks out ~85% of the illuminated area, significantly higher LED intensities were required to induce comparable somatic responses in the proximal stimulation condition.

For PRC measurements, cells were recorded in the cell-attached, current clamp mode (to preclude any feedback from the amplifier that can alter the cells firing pattern). Barrages of light pulses were delivered based on a previously described method [47]. Pulses were 0.5-1 ms long and were separated by random, exponentially distributed inter-pulse intervals, with means of 6 ms and 2.17 ms (from pulse onset until the onset of the next pulse) for the full-field and proximal illumination conditions, respectively. LED light intensity at the back plane of the objective was 15 mW for proximal illumination through the disk with the pinhole and 0.03-0.06 mW for full-field illumination. Stimulation consisted of 3 seconds of baseline recording followed by 9 seconds of barrage stimulation, and this was repeated 25 times. Each of the 25 repetitions was a different realization of exponentially distributed inter-pulse intervals, but all cells received the same set of stimulation barrages. This stimulus was also used for measurements of ChR2 current responses. To avoid the effect of ChR2 deactivation, only data collected during the last 5 seconds of stimulation were used in the analysis.

In electrotonic length experiments, cells were stimulated with a temporal sinusoid at different frequencies (0.25-16 Hz, 3-4 seconds per frequency). LED light intensity at the back plane of the objective was 1.5mW for proximal illumination and 1.5-15 µW for full-field illumination.

In entrainment experiments, cells were recorded in the perforated patch configuration and allowed to fire spontaneously as 30-60 second long sinusoidal waveforms at different frequencies around the natural firing frequency of the cell (typically ~7 Hz) were delivered. Stimulation was applied either optogenetically, in the proximal or full-field illumination configurations, or as a current injection with a 10-20 pA amplitude. LED light intensity at the back plane of the objective was 0.15-0.9 mW for proximal illumination and 15-30 µW for full-field illumination.

### Two-photon laser scanning microscopy (2PLSM)

Neurons were patch clamped with 100 µm Alexa Fluor 568 (Molecular Devices) in the patch pipette. The two-photon excitation source was a Chameleon Vision II tunable Ti:Sapphire pulsed laser (Coherent, Santa Clara, CA, USA) tuned to 820 nm. The images were collected with the Femto2D system (Femtonics, Budapest, Hungary) which includes two 3mm galvo-scanners and a multi-alkaline non-descanned photomultiplier tube for imaging Alexa Fluor. Z stacks of optical sections (spaced 2 µm apart) were collected using 0.2 µm pixels and 15 µs dwell times. The image in Figure 2A is a montage of several collapsed Z stacks. Measurements of the long axis of SNr GABAergic cells somata were based on these images (n=15).

### Data analysis

Data were analyzed and curve fitting was performed using custom made code in MATLAB (MathWorks, Natick, MA, USA).

#### Electrotonic length study

Phase shifts between the optogenetic sinusoidal stimulation and the somatic current response were determined by the location of the peak in the cross-correlation function (CCF) between the two traces, for each stimulation frequency and for each illumination condition. We fit these data with the curve we derive in the Results section (eq. 2) from our cable theoretic model and use the fit to estimate the effective time and characteristic lengths associated with the dendrites of SNr GABAergic neurons. Because of variability in the responses to lower frequency values only values between 6-16 Hz were used to generate the fit.

#### PRC estimation

Steady-state data (4-9 seconds after beginning of stimulation) were analyzed as previously described [47,52], with a few minor changes. In short, spikes were detected and each interspike interval (ISI) was divided into 50 equally-sized bins (i.e., for each ISI, the bin size scales with the duration of the ISI), where the *j*th bin corresponds to the phase Φ_*j*_ = (*j* − 0.5)/50. The number of pulses delivered in each bin of each ISI, denoted *p*_α,*j*_ (α is the index of the ISI) was counted and the mean value (averaged over all ISIs and all bins) was subtracted resulting in δ*p*_α,*j*_. Then, a multiple regression analysis was performed with δ*p*_α,*j*_ values as the independent variables and *ISI*_α_ values as the dependent variables. If we denote the PRC as *Z*(Φ), then the regression coefficients *Z*(Φ_*j*_) provide a unique solution for the PRC [53]. Because the resulting PRCs had a triangular form [47], we fit an acute triangle to the resulting data points. The base of the triangle is the segment from 0 to 1, its peak is located at the phase θ, and its amplitude and offset are *A* and *C*, respectively. A similar triangle with zero offset was used for the analysis of the population responses and entrainment of SNr neurons. As discussed presently, we need the Fourier series of this triangle which is given by 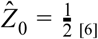 and for all other *k*

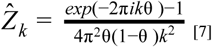

#### Peristimulus time histogram (PSTH) estimation

Because the barrages of pulses are realizations of a time varying Poisson process, with a step-like onset, the same spiking data were used to perform the population rate response analysis to a step-like perturbation. PSTHs centered around the onset of the barrage stimulus were generated. The width of each bin was 4 ms, and data included 25 repetitions of each of the 19 cells. Given the Fourier series of a neuron’s PRC, one can calculate, as shown previously, the predicted shape of the neuron’s PSTH using the Fokker-Planck formalism [54,55]. This calculation yields the following model that we fit to the empirical PSTHs, denoted *R(t)*, corresponding to each of the two stimulation conditions:

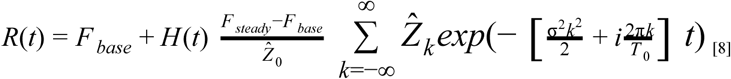

Where *F* _*base*_ is the cells’ average baseline firing rate prior to stimulus onset, *F* _*steady*_ is the average steady-state firing rate after relaxation, *T*_0_ is the baseline period 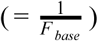, and 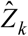 is the *k*th Fouriercoefficient of the PRC (eq.6 and eq.7). *H(t)* is the Heaviside function.

We fit *R(t)* (eq. 8) to the observed baseline and steady-state rates of the PSTH and the structure of the first 50 ms after stimulus onset. The parameters that were estimated were σ and θ, for each stimulation condition, and we used *k* values ranging from −100 to 100.

For ChR2 current response measurements, a double exponent of the form 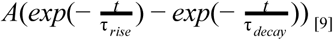 was fit to average current response curves, and the parameter *A* as well as the time constants *τ* _*rise*_ were τ _*decay*_ extracted [47]

#### Entrainment to rhythmic inputs

*Entrainment of the cell’s spiking activity to rhythmic inputs was evaluated based on previously described methods [57]. For each spike, the perturbed period T*_*p*_ was measured as a function of its effective phase Ψ - its phase with respect to the period of the rhythmic stimulus. The fitted curve for these measurements (Fourier fit, first 3 modes) was used to generate an iterative map of the effective phase Ψ_*n*+1_ of the next action potential as a function of the effective phase Ψ_*n*_ of the current action potential (see eq. 3 in Results). Cobweb plotting was applied to this map to investigate the dynamics of the evolution of effective phases [57]. In order to generate a prediction for phases of locking using proximal and full-field illumination, a differential equation describing the evolution of the cell’s intrinsic phase (see eq. 5 in Results) was numerically integrated in XPPAUT [80], using the idealized triangular PRC fit, with the values of θ extracted from the estimate of the PSTH.

The probability distribution of effective phases was used to generate a circular variance vector, which provides a measure of the variation of effective phases [60]. The amplitude of the circular variance vector represents the extent to which the firing of the neuron was entrained by the oscillatory input, and its phase indicates the effective phase of entrainment, which corresponds to the peak of the distribution of effective phases. Each cell was stimulated with temporal cosine waveforms at several different frequencies (2.5-21 Hz), and a circular variance vector was estimated for each trial. In order to investigate phases of entrainment, we only included instances where the amplitude of the circular variance was larger than 0.175. This threshold represents the 95th percentile in the distribution of the amplitude of circular variances achieved by bootstrapping. The surrogate data were series of 100 uniformly distributed random numbers (between 0 and 1), because our entrainment experiments typically included on the order of 100 ISIs.

#### Statistics

The nonparametric two-tailed Wilcoxon rank-sum test (RST) was used for non-matched samples, and that nonparametric Wilcoxon signed-rank test (SRT) was used for matched samples. Error bars represent standard errors of the mean. The parametric ANCOVA test was used to test significant changes in curves. In PRC analysis the test was applied to the rise stage of the curves.

In the entrainment analysis, P-values were derived by bootstrapping. To test whether the distribution of proximal circular variance vectors is significantly different from the distributions induced by somatic current injection and full-field illumination, we calculated the probability to get the empirical average phase obtained in the latter two conditions assuming that samples came from the proximal condition distribution. To make the problem one dimensional, the representation of each phase in the proximal condition distribution was determined by the size of the vector. Current injection and full-field illumination distributions were compared by repeating this process with current injection phases serving as the surrogate data.

Null hypotheses were rejected if the P-value was below 0.05.

## Supporting information

## Acknowledgments

We would like to thank Dr. Charles J. Wilson for illuminating discussions about this study. Eng. Anatoly Shapochnikov provided excellent technical support. This work was funded by a European Research Council (ERC) Consolidator Grant (no. 646886) to J.A.G.

## Author Contribution

L.T. conceptualized, designed and conducted the experiments, analyzed the data, developed the theory and conducted simulations and wrote the paper. J.A.G. conceptualized, designed and supervised the experiments, developed the theory and conducted simulations, wrote the paper, secured funding and is responsible for the content of the manuscript.

