## Supplementary material for "Population dynamics and entrainment of basal ganglia pacemakers are shaped by their dendritic arbors"

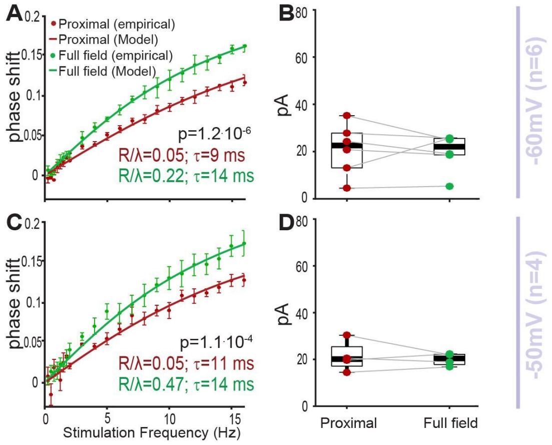


**S1 Figure: Shift in somatic current responses to periodic illuminations of SNr GABAergic neurons at various holding voltages.** Channelrhodopsin-2-expressing SNr neurons were optogenetically stimulated using sinusoidally modulated light at various temporal frequencies, while blocking all inputs to the cell. The cell was voltage clamped at –60 mV (*top row*) or -50 mV (*bottom row*), and the current injected to the soma was measured in the whole-cell configuration. **A:** When the cells are held at -60 mV, proximal and full-field illumination conditions give rise to two distinct curves (n=6, $p=1.2 \cdot 10^{-6}$; ANCOVA). The response to rhythmic stimulation exhibits a spatial effect – phase shift between stimulation and somatic response is larger under full-field illumination (green) compared to proximal (red). Error bars represent standard error of the mean. **B:** Amplitude of somatic current response was comparable between the proximal and full-field stimulation conditions. Central marks indicate the median and box edges represent 25th and 75th percentiles. **C:** Same as A with cells held at -50 mV (n=4, $p=1.1 \cdot 10^{-4}$; ANCOVA). **D:** Same as B with cells held at -50 mV.
