## Supplementary material for "Population dynamics and entrainment of basal ganglia pacemakers are shaped by their dendritic arbors"

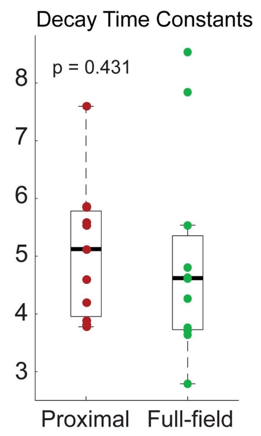


**S2 Figure: Current response to pulse illumination in the full-field configuration is slower to rise than in the proximal configuration.** For each stimulation condition, a double exponential was fit to the average current responses for each cell (see Methods and Materials) and rise and decay time constants were extracted ( $\tau_{rise}$ , $\tau_{decay}$ ). The distributions of decay time constants are not significantly different. Central marks indicate the median and box edges represent 25th and 75th percentiles.
