## Supplementary material for "Population dynamics and entrainment of basal ganglia pacemakers are shaped by their dendritic arbors"

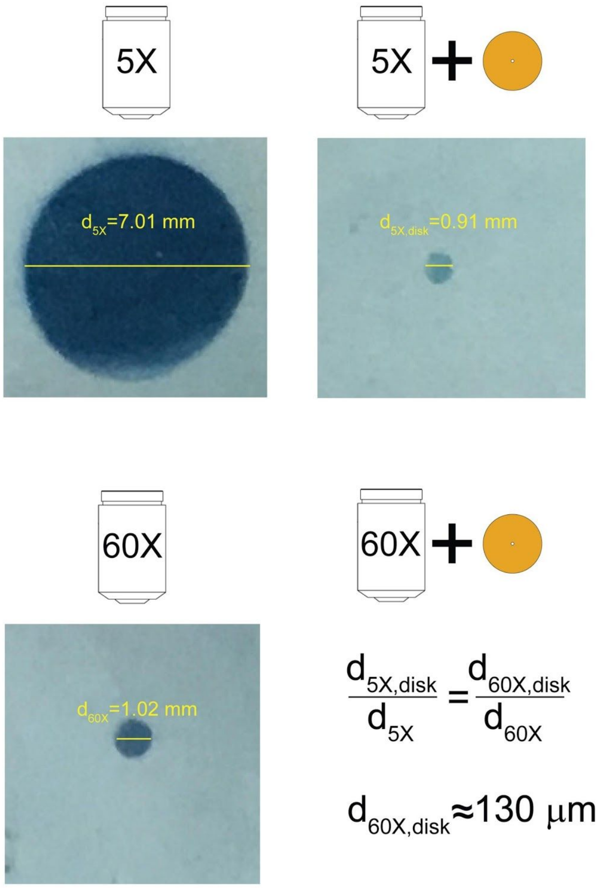


**S3 Figure: Measurements of diameters of illuminated regions for proximal and full-field stimulation conditions.** Measurements were done by placing a light sensitive paper (SunArt Paper, TEDCO toys, Washington, IN) which changes its color when exposed to light at the focal plane of the relevant objective and measuring the diameter of the discolored region. We used two stimulation conditions: 1) proximal stimulation achieved by illuminating through a water-immersion 60X objective and placing an opaque disk with a pinhole at the objective's rear focal plane (*bottom right*); and 2) full-field stimulation achieved by illuminating using a 5X objective (*top left*). When full-field illumination was used, the diameter of the discolored area was $d_{5X} \approx 7\ mm$. In the case of the proximal stimulation condition, the diameter $d_{60X,disk}$ could not be measured reliably using this method because the affected area is very small and the light is very weak. Instead, we measured the diameter $d_{5X,disk}$ that corresponds to illuminating through a 5X objective with the pinholed opaque disk placed at the objective's rear focal plane (*top right*). The measured diameter was $d_{5X,disk} \approx 0.91\ mm$, and we concluded that the disk restricts the illuminated area making the new diameter 13% of the original diameter when the disk is not used. Next, we measured the diameter of the illuminated area when a 60X objective is used without the pinholed disk (*bottom left*). We used the measured diameter $d_{60X} \approx 1\ mm$ to estimate the diameter of the activated region under the proximal stimulation condition (*bottom right*): $d_{60X,disk} = \frac{d_{5X,disk}}{d_{5X}} \cdot d_{60X} \approx 130\ \mu m$. Thus, the diameters of stimulated regions in the proximal and full-field stimulation conditions are $130\ \mu m$ and 7 mm, respectively. Because water was used to develop the light-sensitive paper, the $d_{60X}$ diameter was measured with the lens immersed in ethanol that has a refractive index (1.36) that is very close to that of the ACSF which is water with a high salt content.
